## Supplemental Information for "Existing function in primary visual cortex is not perturbed by new skill acquisition of a non-matched sensory task"

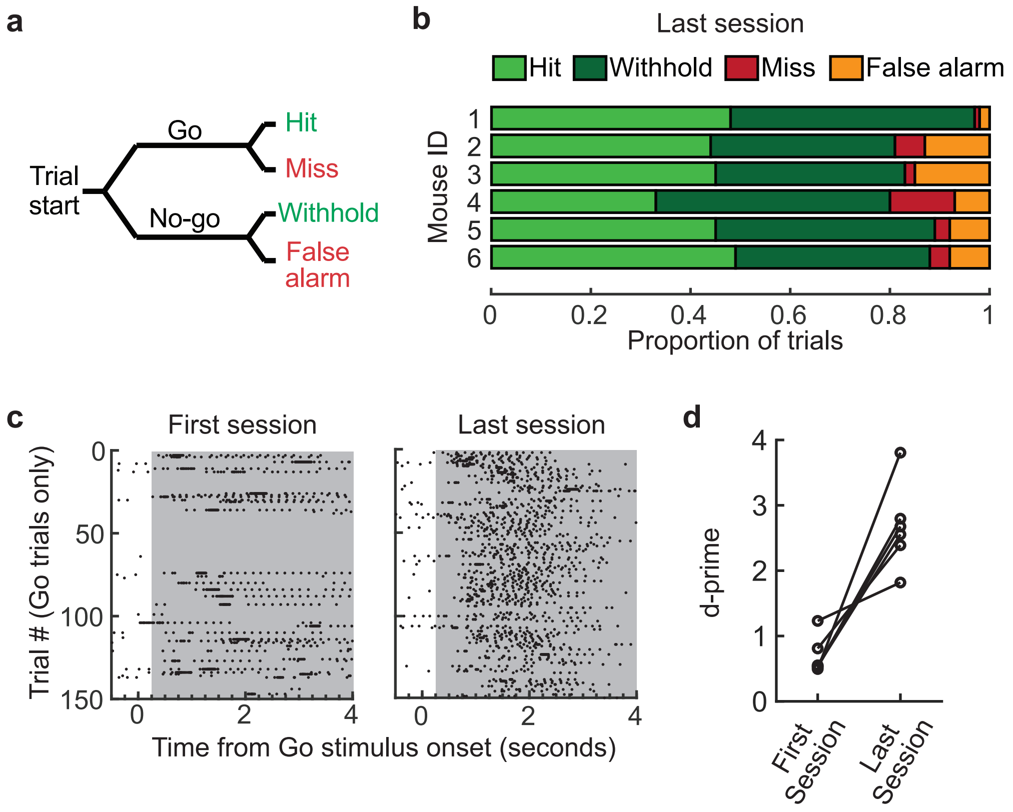

**Supplementary Figure 1. Mice learned to associate a 15 kHz pitch with reward prior to BCI training**

(a) Schematic depicting four possible trial outcomes of the Go/No-go auditory discrimination shaping task. The ‘Go’ stimulus was a 15 kHz pitch and ‘No-go’ stimulus was a 5 kHz pitch. Mice performed 200 to 500 trials in one session per day; one mouse was removed because it consistently failed to perform at least 50 trials per session. It is our experience that a minimum of 75 trials per session is required for mice to learn associations in Go/No-tasks.

(b) The proportion of trial outcomes for 6 mice on the last session. Session performance is defined as the highest proportion of correct behavioral responses (hits and withholds, shown as green and dark green) calculated over a sliding window of 100 consecutive trials. Six of 7 mice trained on this task reached the 80% performance criterion and moved on to BCI training or BCI control experiments. Days to reach criterion in parentheses: mouse #1, 97% (5 days); mouse #2, 81% (16 days); mouse #3, 83% (12 days); mouse #4, 80% (8 days); mouse #5, 89% (5 days), or BCI control: mouse #6, 88% (6 days).

(c) Raster of lick times aligned to the onset of the ‘Go’ stimulus for an example mouse on the first and last session. Reward became available after a 250 ms delay from the onset of the ‘Go’ stimulus, if the animal licked, and remained available for the duration of stimulus presentation (gray). Mice were free to lick more than once on ‘Go’ trials but received only one water drop per trial. Note, there are more missed trials on the first session (114 missed trials out of 150 ‘Go’ trials) compared to the last session (27 out of 149).

(d) Discrimination accuracy of 6 mice in the first and last session reported as d-prime, computed from 100 trials. All 6 mice had a d-prime of at least 1.68 on the last session.

Source data are provided as a Source Data file.

S2

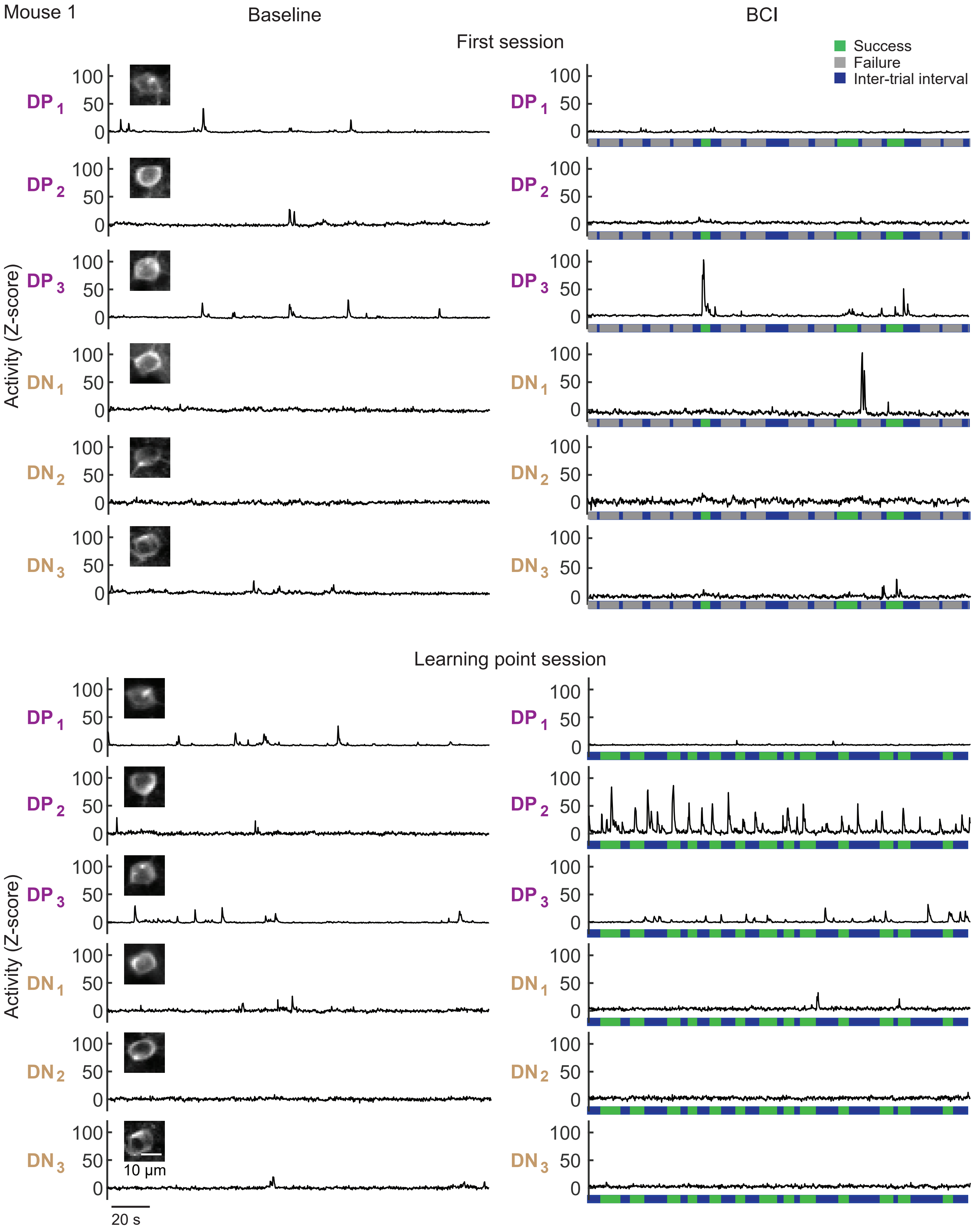

**Supplementary Figure 2. Examples of direct neuron activity, Mouse 1.**

Example activity of the 6 direct neurons (200 ms for each condition). Data are re-plotted from Fig. 1c and include y-axis values to demonstrate that the signal was stable within a given session, and across sessions. Consistent with the idea that cross-session changes in baseline activity did not drive improved performance, cases in which individual DP neurons exhibited high event rates during BCI but not baseline were observed (DP neuron #2 in mouse #1, for example). We noted that within a given mouse, the height of sample-to-sample fluctuations varied (see mouse # 3 for example, DP neuron #1 had smaller fluctuations than DP neuron #2, Fig. S4). Smaller sample-to-sample fluctuations are expected in neurons with higher basal firing rate, such as exhibited by DP neuron #1 in mouse #3. The same Y-axis scale was applied to all neurons within a given animal to facilitate within-animal, cross-session comparisons.

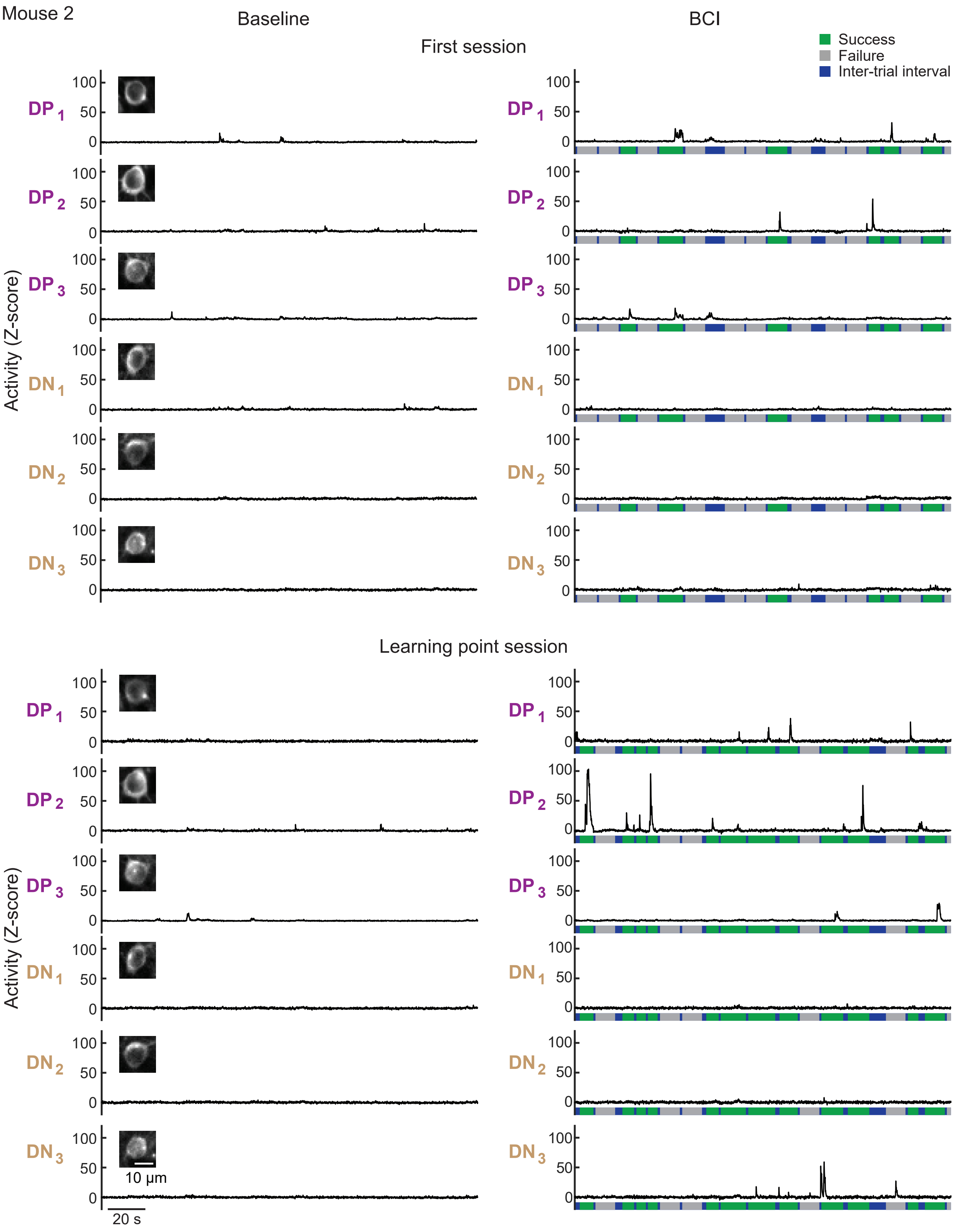

**Supplementary Figure 3. Examples of direct neuron activity, Mouse 2.**
Example activity of the 6 direct neurons. Labels as in Supplementary Fig. 2.

Mouse 3

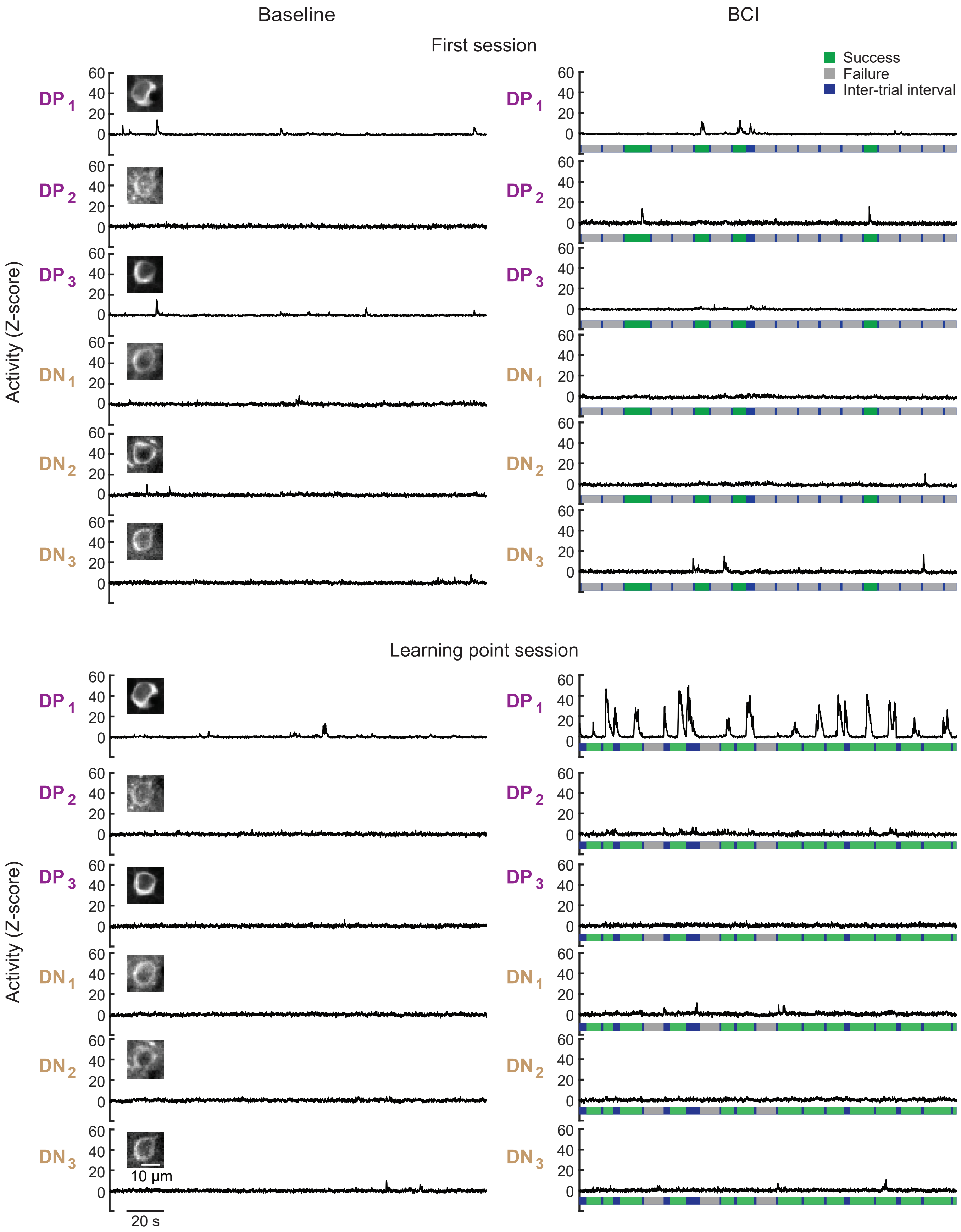

**Supplementary Figure 4. Examples of direct neuron activity, Mouse 3.**
Example activity of the 6 direct neurons. Labels as in Supplementary Fig. 2.

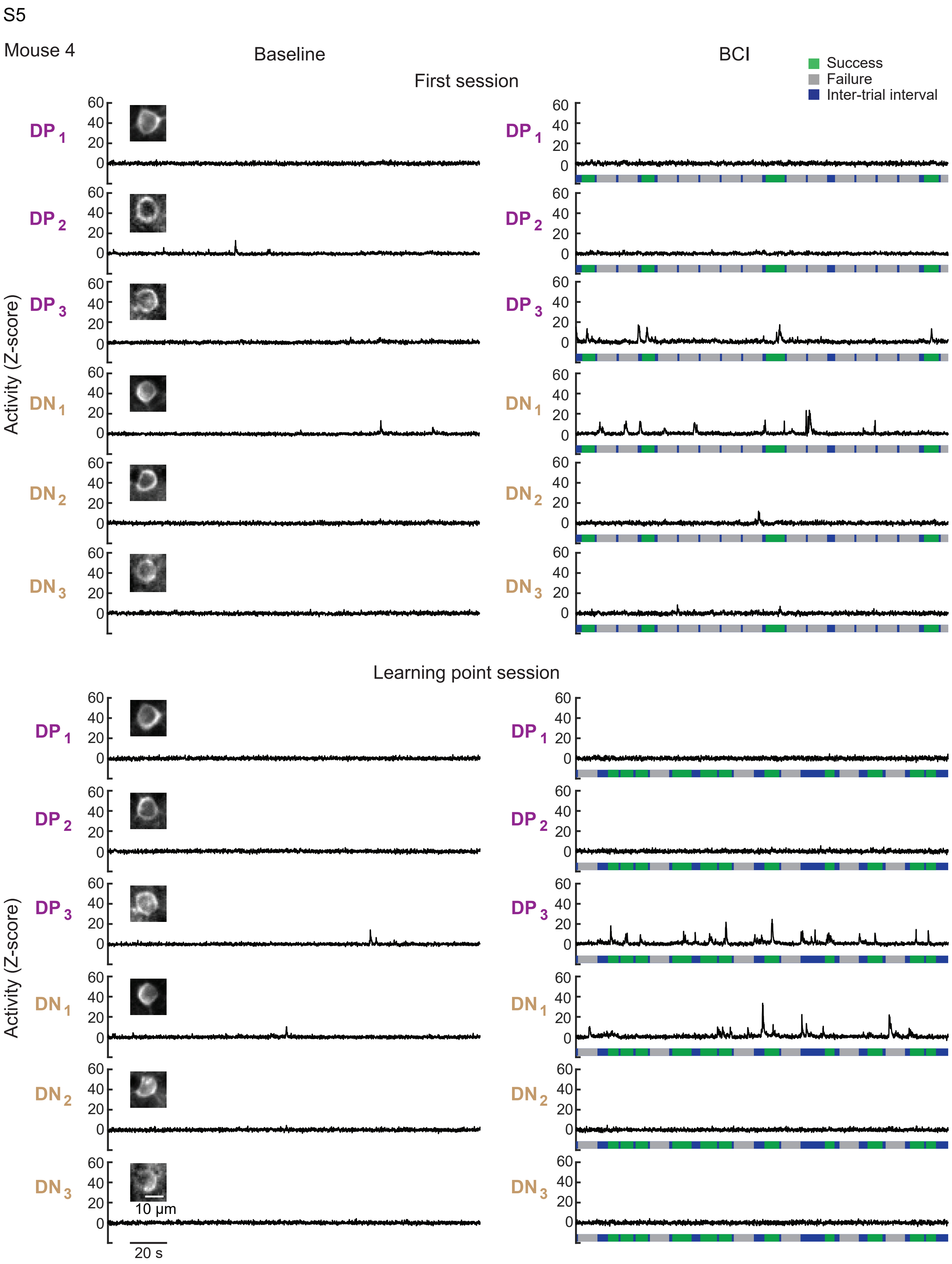

**Supplementary Figure 5. Examples of direct neuron activity, Mouse 4.**
Example activity of the 6 direct neurons. Labels as in Supplementary Fig. 2.

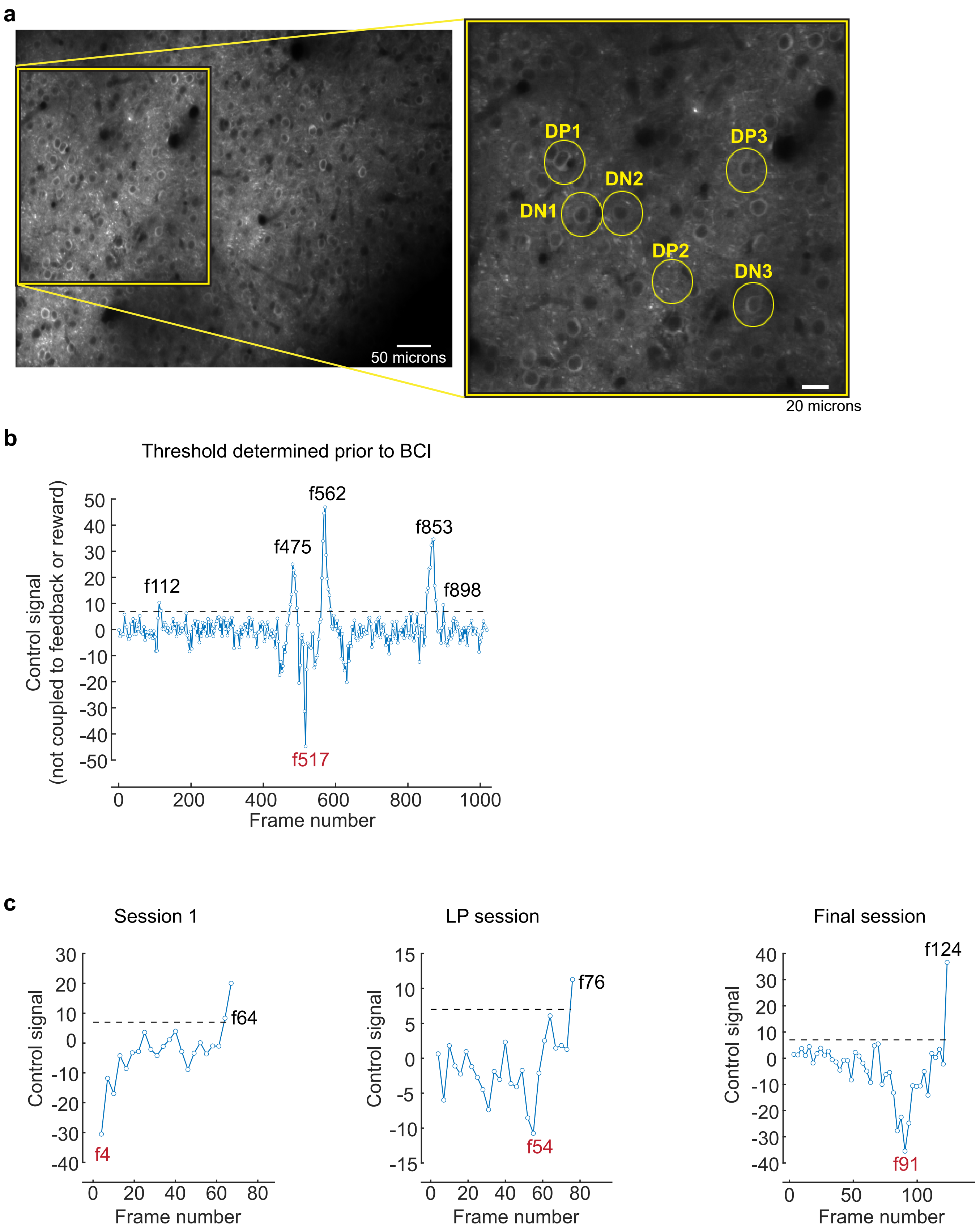

**Supplementary Figure 6. Selection of direct neurons.**

(a) Spatial location of the 6 direct neurons selected in mouse #3.

(b) Control signal target threshold crossings (dashed line; imaging frame indicated, f). Note, the control signal is summed across three frames (circles). The maximum negative drive is indicated in red. See Supplementary Movie 1.

(c) Example trials within a given session as indicated, labels as in 'b'. See Supplementary Movies 2-4.

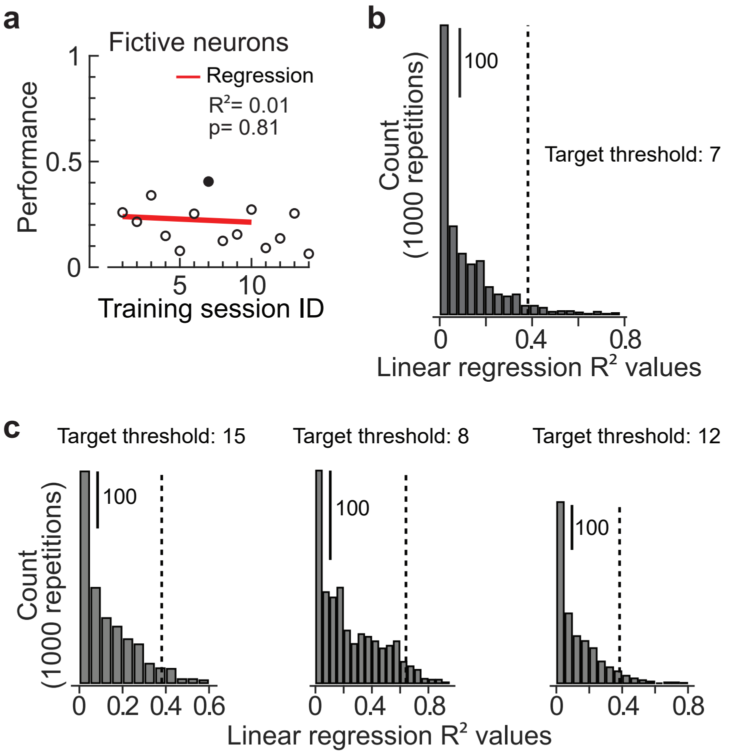

**Supplementary Figure 7. Performance does not spontaneously increase in an auditory playback control mouse.**

In the auditory playback control mouse, no actual neurons were coupled to the auditory pitch or reward delivery, and the auditory feedback generated by mouse #1 on sessions 1-10 was played back to this control mouse. Six 'fictive' direct neurons were selected using the same procedure as in the BCI mice (the threshold was 7 in this case), and tracked across 14 sessions. For sessions 11-14 in the control mouse, the trial order of the feedback generated by mouse #1 on session 10 was randomized and played back. Water reward was available in response to licking when the auditory signal reached 15 kHz to ensure that the control mouse was engaged in the task. In all sessions the control mouse licked on the 15 kHz tone in 82% or more of the trials.

(a) Performance of the fictive direct neurons did not improve over the course of the 14 sessions, and remained below 0.5 for all sessions. One out of 14 sessions scored as having a significantly higher performance than session 1 (black, 2-sample proportions Z-test, corrected for 14 multiple comparisons). Linear regression was calculated across sessions 1-10; in contrast to the 4 BCI mice, it was not significant ($p = 0.81$, $n = 10$ sessions).

(b) The null distribution ($n= 1000$) of $R^2$ values calculated from indirect neurons, excluding the 6 fictive neurons, using sessions 1-10 and a threshold of 7 is shown. The 95$^{th}$ percentile of this distribution (one-sided) is 0.38 (dashed line).

(c) In addition to a target threshold of 7, the null distributions ($n= 1000$) of $R^2$ values were calculated as in 'b' for the other three target thresholds used in this study. The dashed lines indicate the 95$^{th}$ percentile of the distribution. In all cases the actual performance of the BCI mouse was higher than the 95$^{th}$ percentile of its matching null distribution (one-sided).

Source data are provided as a Source Data file.

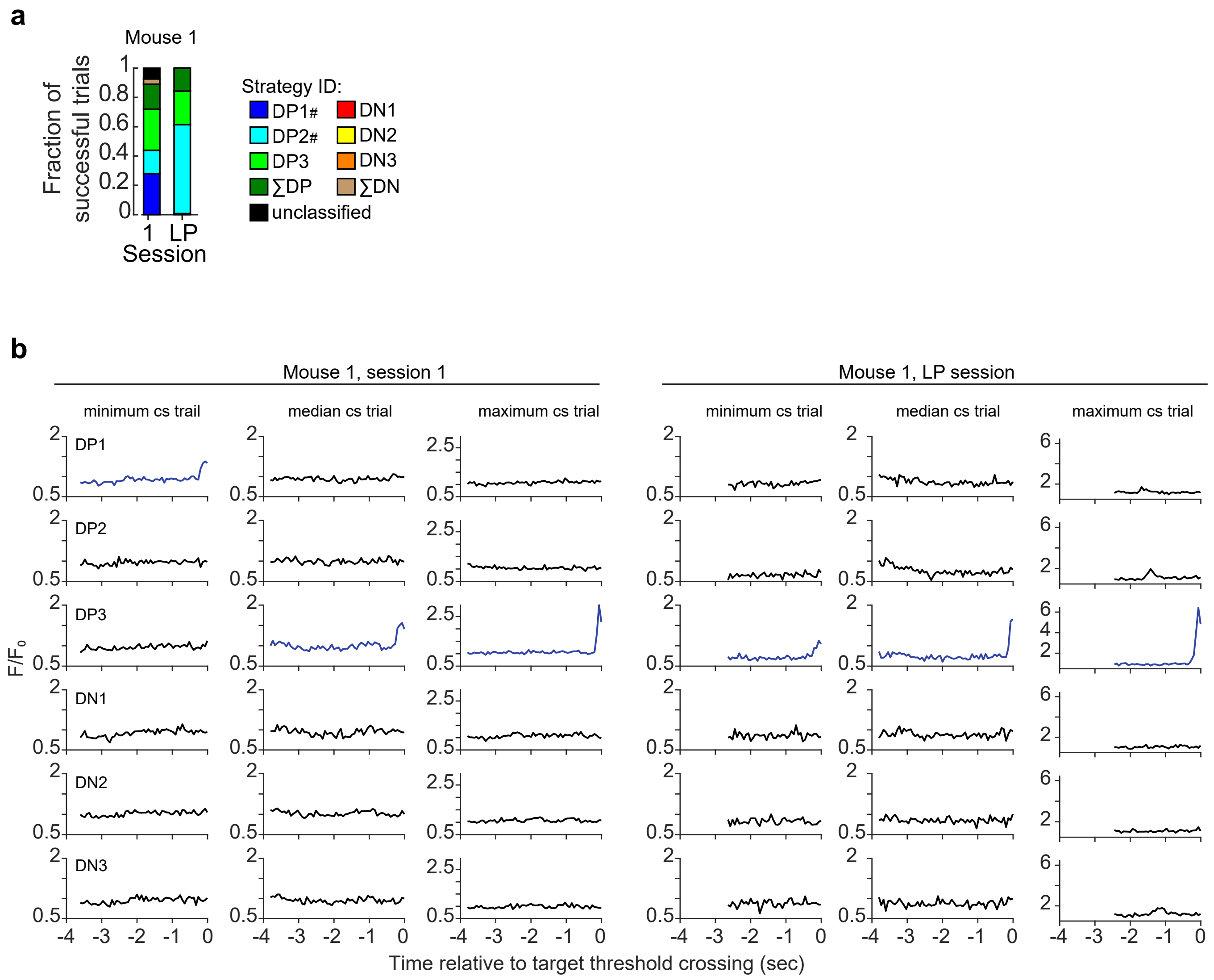

**Supplementary Figure 8. Example trials of direct neuron activity, Mouse 1.**

(a) Strategy categories, defined in Fig. 2 (replotted here for easy reference). DP neurons that significantly changed (2-sample proportions Z-test, corrected for 3 multiple comparisons) their contribution to threshold crossings (session 1 versus the LP session) are indicated (#).

(b) The BCI trial with the minimum amplitude of control signal (CS) was identified and the fluorescent activity of the individual direct neurons plotted here, aligned to target threshold crossing. Direct neurons DP 1-3 and DN 1-3 are indicated, by row. The direct neurons that contributed to driving the auditory cursor closer to target threshold at the time of success are indicated in blue. The trial with the median and maximum control signal as shown for comparison.

Source data are provided as a Source Data file.

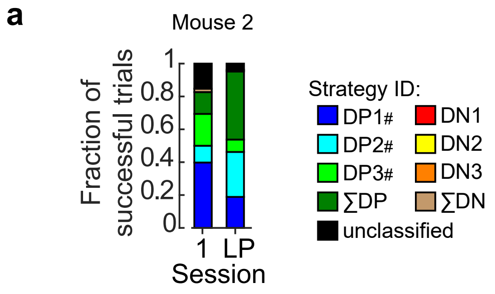

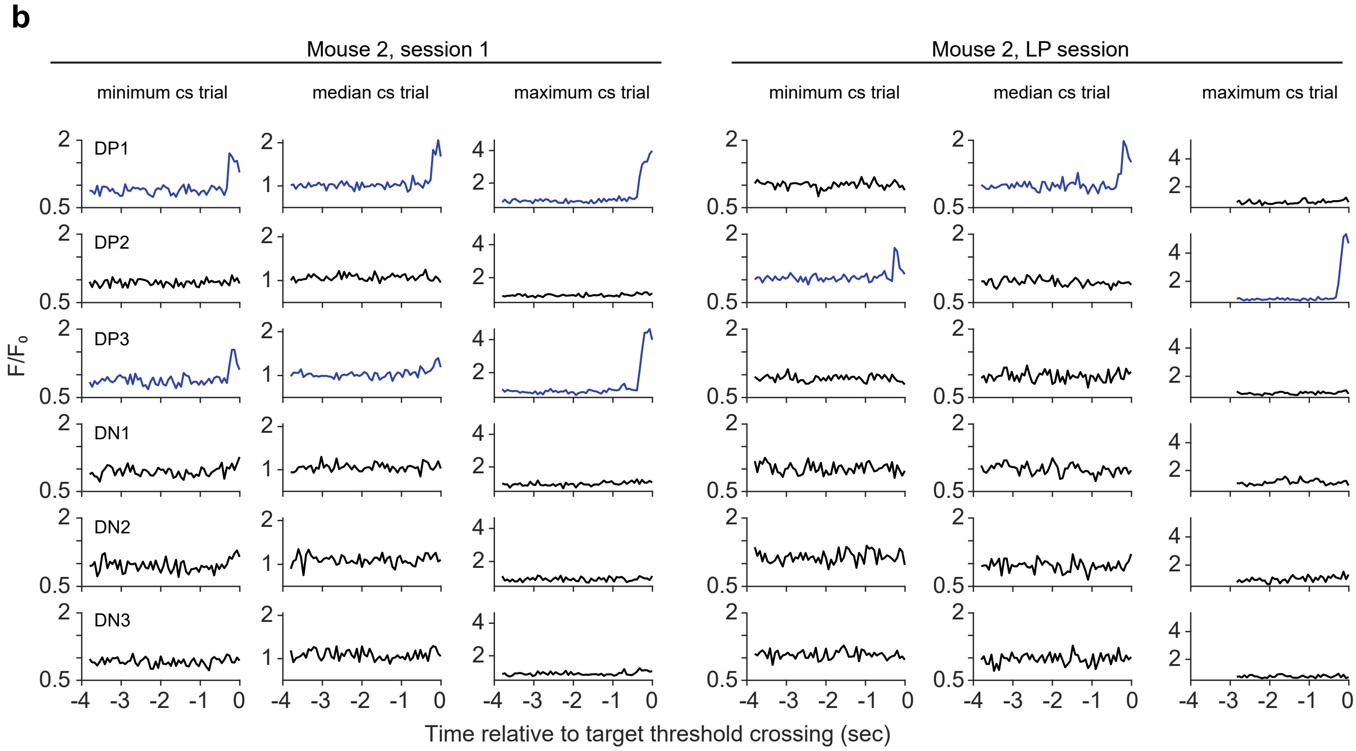

**Supplementary Figure 9. Example trials of direct neuron activity, Mouse 2.**

(a) Strategy categories, labels as in Fig. 2.

(b) BCI trials with the minimum, median, and maximum amplitude of control signal are shown. Labels as in Supplementary Fig. 8.

Source data are provided as a Source Data file.

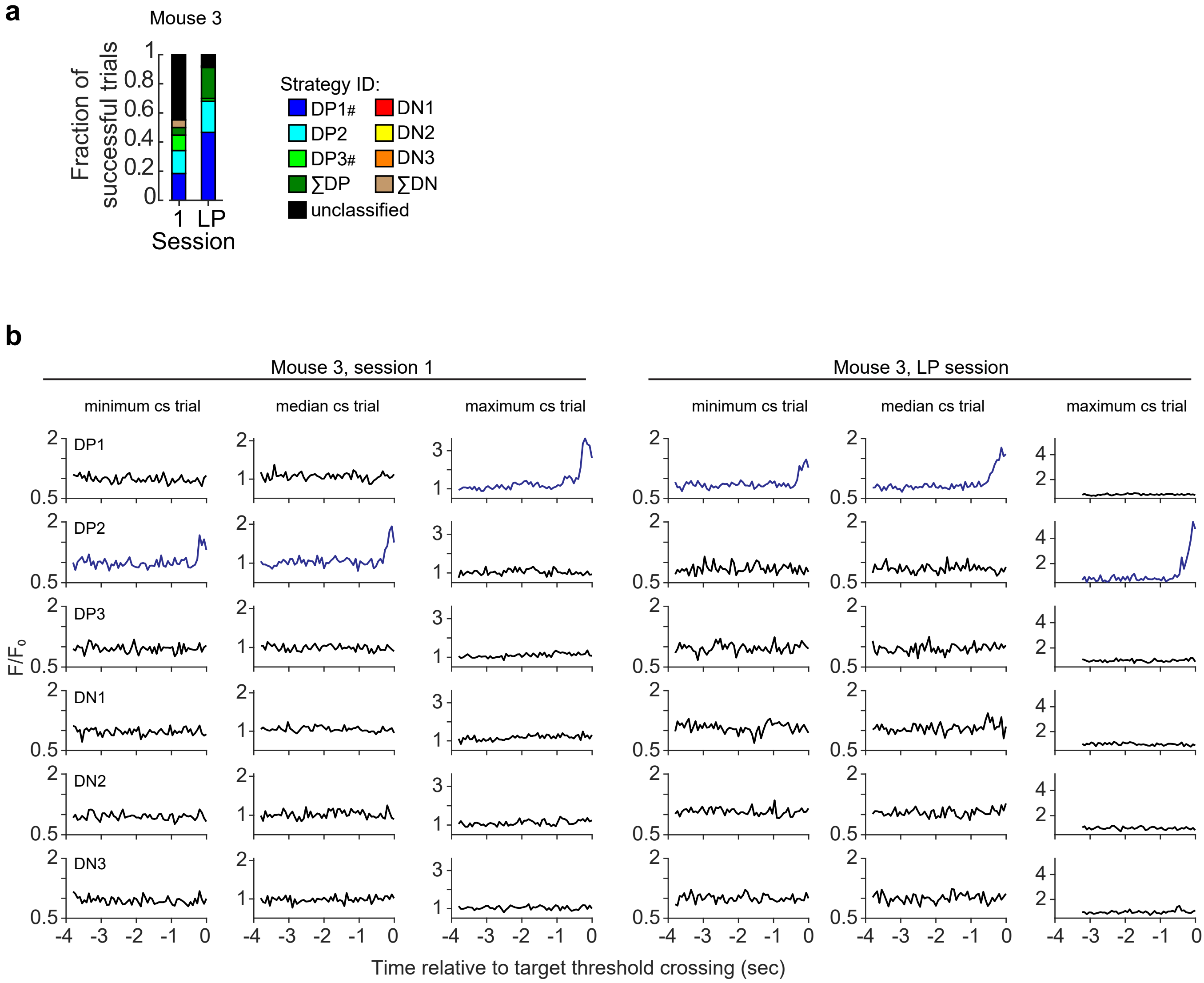

**Supplementary Figure 10. Example trials of direct neuron activity, Mouse 3.**

(a) Strategy categories, defined in Fig. 2.

(b) BCI trials with the minimum, median, and maximum amplitude of control signal are shown. Labels as in Supplementary Fig. 8.

Source data are provided as a Source Data file.

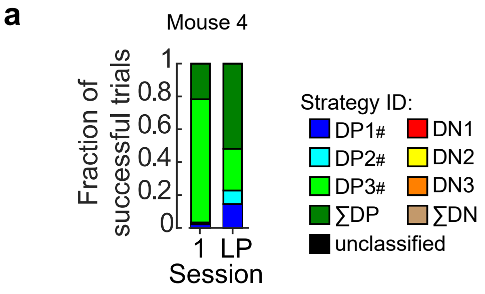

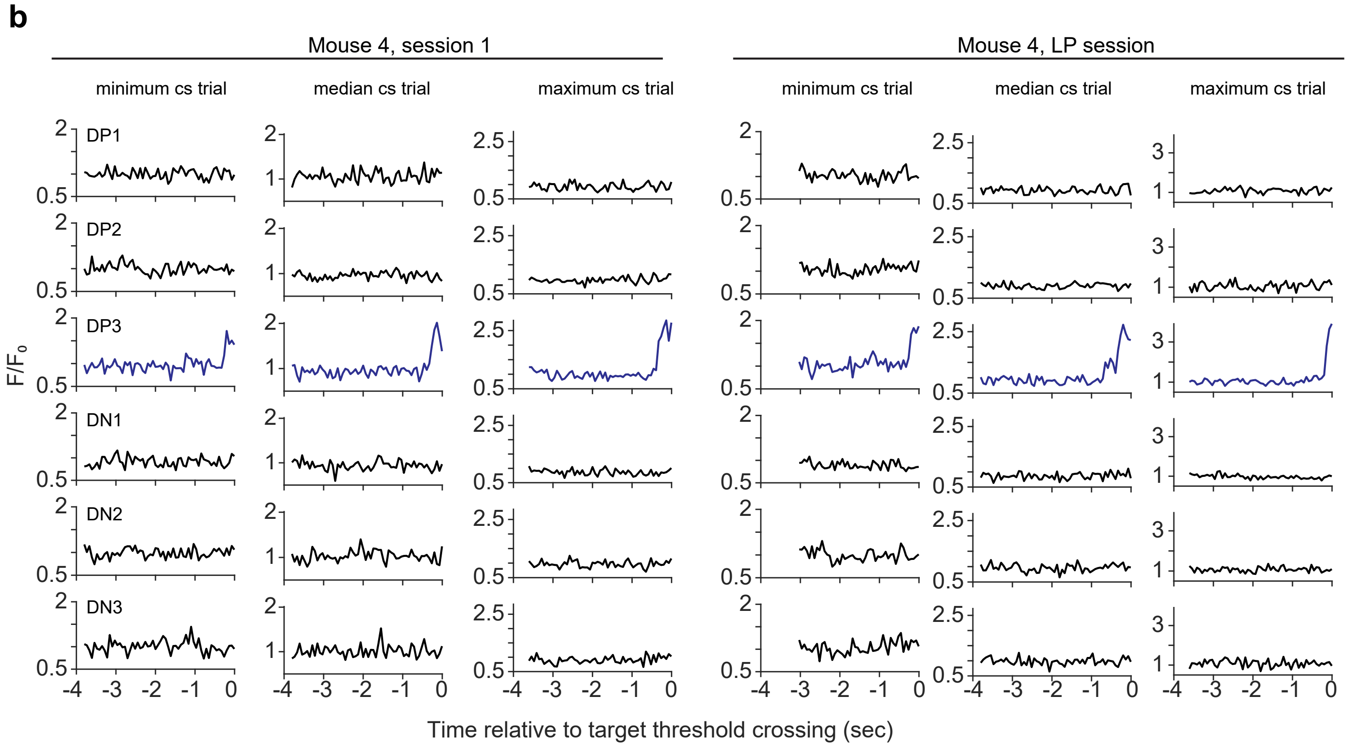

**Supplementary Figure 11. Example trials of direct neuron activity, Mouse 4.**

(a) Strategy categories, defined in Fig. 2.

(b) BCI trials with the minimum, median, and maximum amplitude of control signal are shown. Labels as in Supplementary Fig. 8.

Source data are provided as a Source Data file.

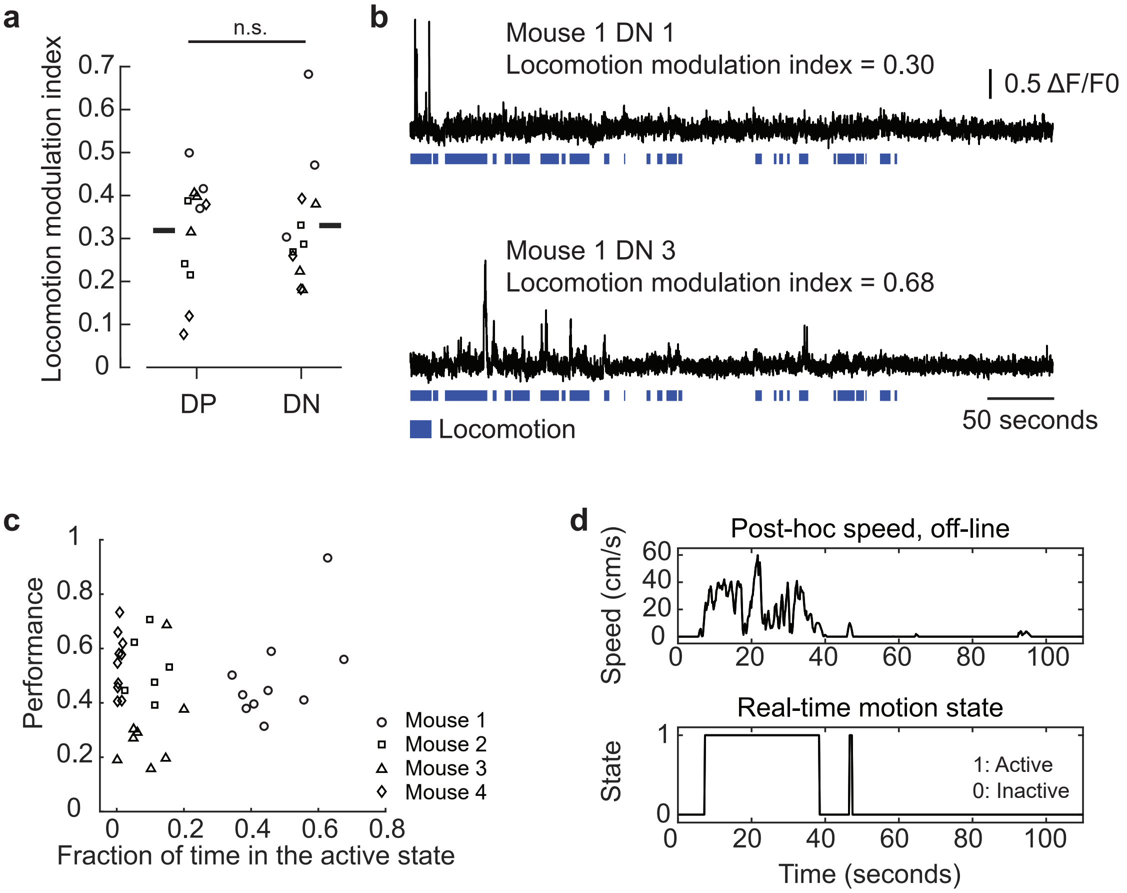

**Supplementary Figure 12. BCI performance was not correlated with locomotion.**

(a) Locomotion modulation index of direct neurons. There was no difference detected between DP and DN neurons (Wilcoxon rank-sum test, $p = 0.751$, $n = 12$ neurons for each group). Horizontal bars represent the average across all DP and DN neurons, respectively

(b) Fluorescence traces of two example direct negative neurons; bouts of locomotion are highlighted (blue).

(c) BCI session performance plotted as a function of the duration of time spent in an active locomotor state. Pearson correlation p= 0.432, n = 34 sessions. Symbols are the same as in 'a'.

(d) Demonstration of how the duration of active states was computed. During BCI experiments, using the tracking laser sensor, locomotion speeds greater than 7 cm/s defined the onset of individual activity bouts, and the offset of each activity bout was defined as the point in time when the locomotion speed was less than 7 cm/s for at least 2 seconds. (bottom). Top, off-line locomotion speed, acquired with a camera. See Supplementary Movie 5.

Source data are provided as a Source Data file.

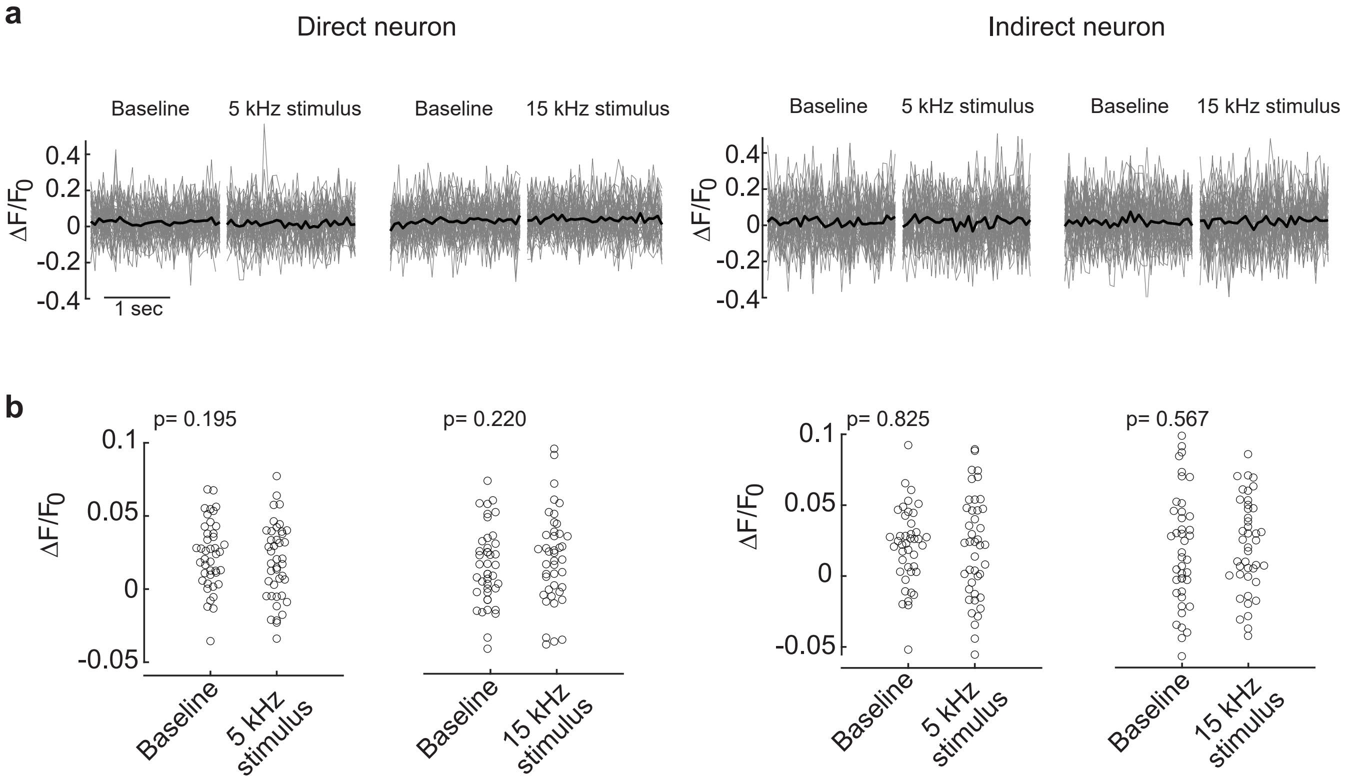

**Supplementary Figure 13. V1 neurons were not modulated by auditory pitches experienced during BCI training.**

After BCI learning (8±6 days after the learning point session), calcium imaging was performed in 3 of 4 mice while mice were presented the 6 auditory pitches used during BCI training. Auditory pitches were presented at least 30 times in a random order for 2 seconds, and each presentation was interleaved with 2.5 seconds of silence; the 2 seconds preceding stimulus onset is defined as baseline ($F_0$). To test for modulation, the activity of a given neuron during the 2 second stimulus presentation was compared to the baseline. Baseline for a given neuron was calculated by averaging the activity across all baseline periods.

(a) Example activity during baseline and sound stimulus, one direct neuron (42 trials, gray) and one indirect neuron (42 trials, gray). The mean across all trials is shown in black.

(b) No difference in activity between baseline and stimulus epochs was detected (paired t-test, 5 kHz $p = 0.195$; 15 kHz, $p = 0.220$, $n = 42$ trials for each stimulus) for the example neurons shown in 'a'. Across the three mice, less than 2% (9 out 495) of all neurons had a mean difference between baseline and stimulus greater than 0.1 $\Delta F/F_0$. Of those neurons, none were significantly modulated by any of the 6 auditory pitch stimuli (paired t-test, $\alpha=0.05$).

Source data are provided as a Source Data file.

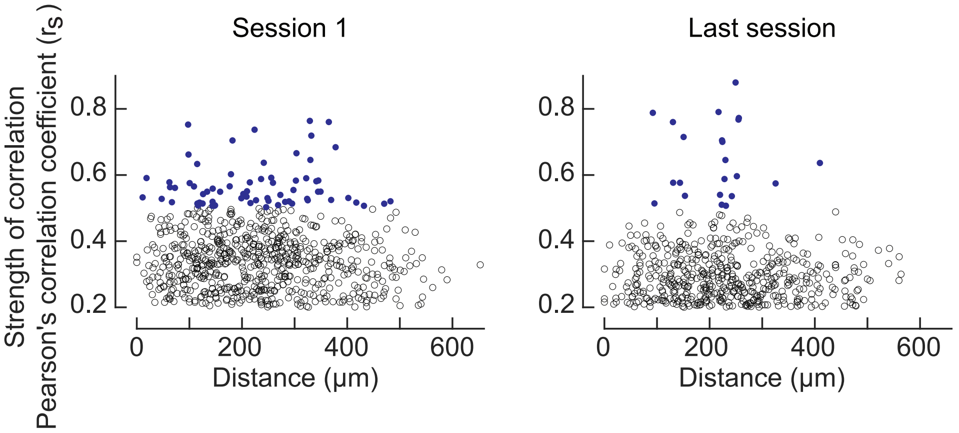

**Supplementary Figure 14. Indirect-DP pairwise correlations were not dependent on distance.**
Distribution of all indirect-DP neuron pairs that were significantly correlated (Pearson's correlation, α=0.05) at the time of success, across trials for the first and last BCI sessions, pooled across mice. There was no relationship between the strength of indirect-DP correlation ($r_s$) for a given pair and the distance between that pair in either the first or last BCI session (Pearson's correlation, r= -0.07, p=0.067, n = 733 pairs and r= -0.03, p=0.465, n = 606 pairs respectively). Similarly, the spatial position of indirect-DP neuron pairs with a magnitude of correlation >0.5 (blue data points) were also not correlated with distance in the first or last session (Pearson's correlation, r=0.08, p= 0.503, n = 70 pairs and r=0.01, p= 0.961, n = 22 pairs respectively).
Source data are provided as a Source Data file.

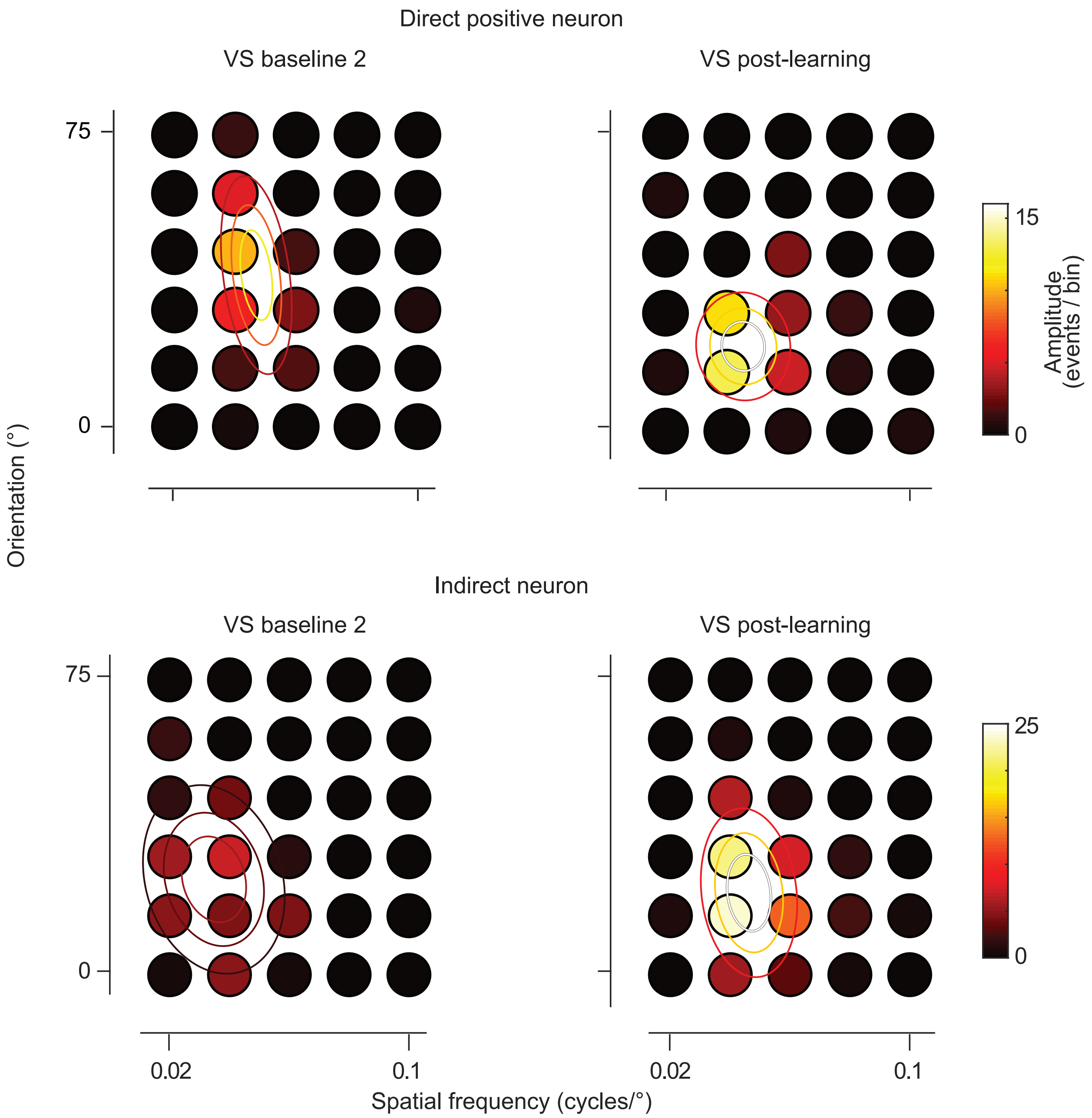

**Supplement Figure 15. Example 2-dimensional tuning profile, cropped.**
Data are re-plotted from Fig. 4b using a smaller range of orientations and spatial frequencies.

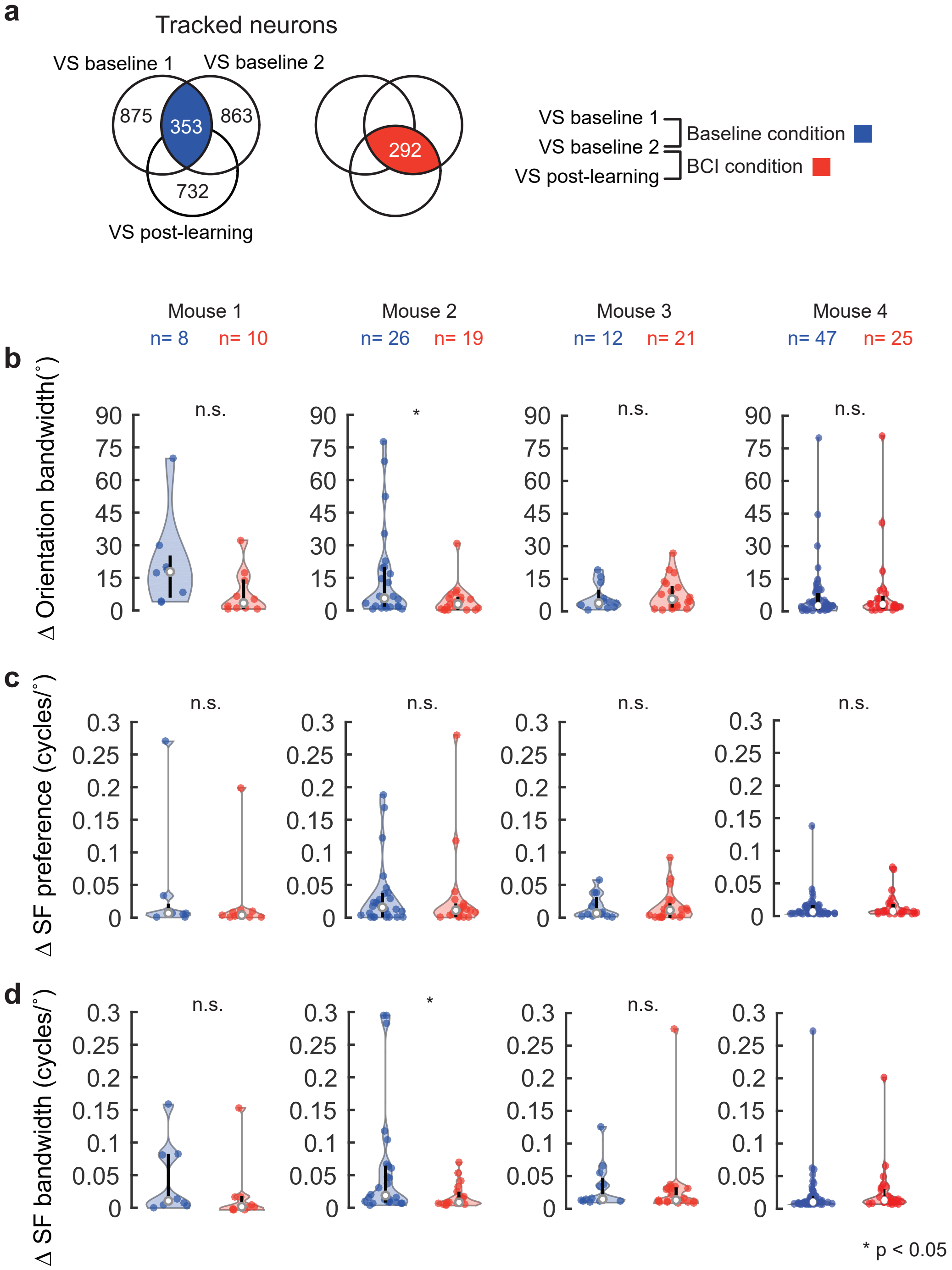

**Supplementary Figure 16. Orientation and spatial frequency tuning parameters remain stable after BCI learning.**

(a) Venn diagrams, re-plotted from Fig. 5a.

(b-d) Stability of tuning parameters in the baseline condition (blue) and BCI condition (red) as indicated. Violin plot labels as in Fig. 4d. Distributions for the three parameters were overlapping in all 4 mice. Wilcoxon rank-sum test p values (mouse #1-4, respectively): Orientation bandwidth, p=0.055, p=0.042, p=0.667, p=0.869. Spatial frequency preference, p = 0.573, p = 0.573, p = 0.587, p = 0.653. Spatial frequency bandwidth, p=0.122, p=0.038, p=0.270, p=0.189. Number of neurons in baseline and BCI condition for mouse #1-4: n=8, 10; n=26, 19; n=12, 21; n=47, 25).

Source data are provided as a Source Data file.

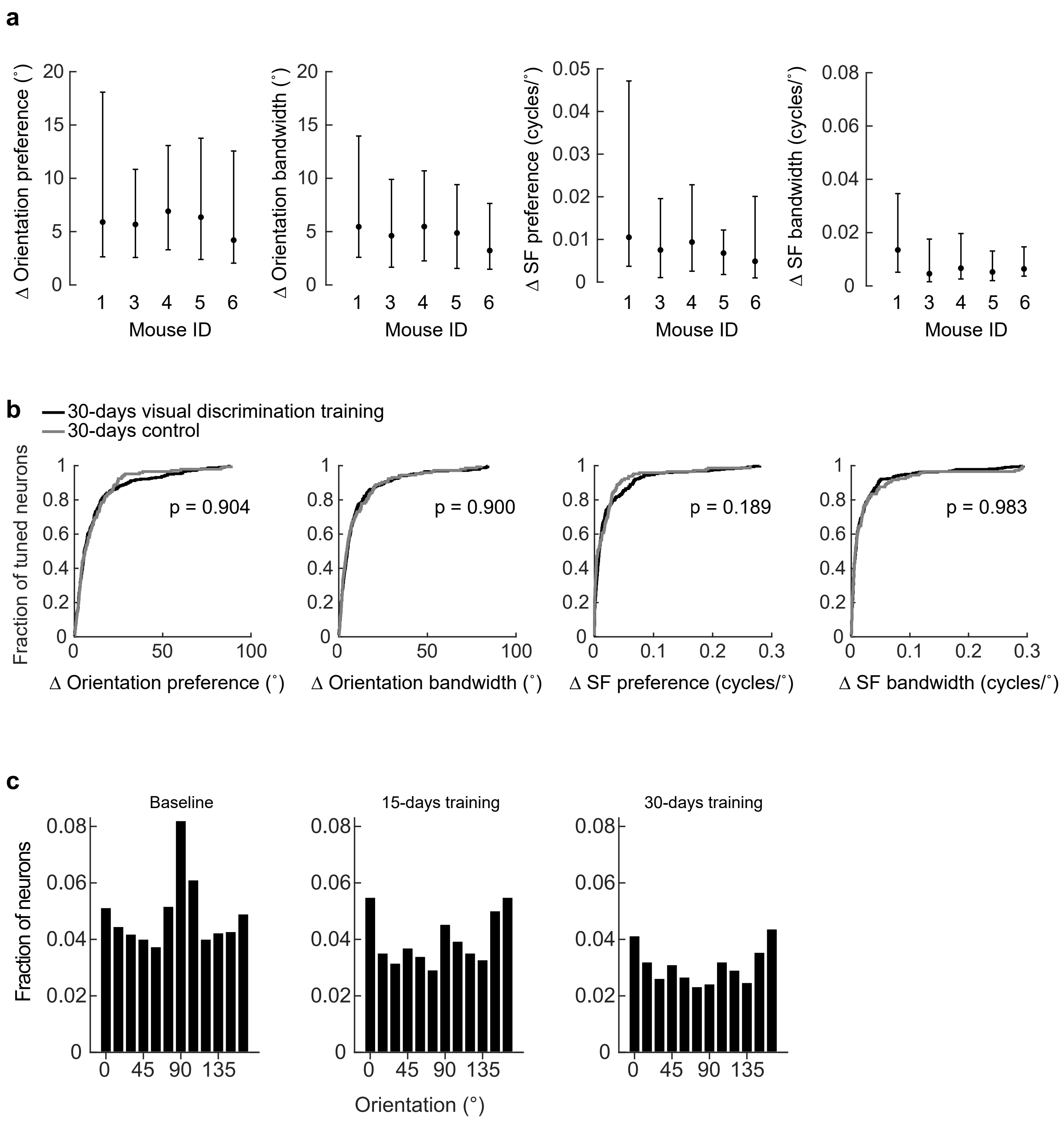

**Supplementary Figure 17. Orientation and spatial frequency tuning parameters remain stable after visual discrimination learning.**

(a) Medians and the interquartile range of the change (Δ) in 4 tuning parameters, as indicated, after 30 days of visual discrimination training for each of the 5 mice shown in Fig. 6c ( mouse #1-5, respectively: 90,56,86,43, and 45 neurons).

(b) Distribution of the change (Δ) in 4 tuning parameters, as indicated, in 30-day control and visual discrimination conditions (K-S test, p-values for the 4 tuning parameters: p=0.904, p=0.900, p=0.189, p=0.983, control n=145 tracked neurons, 5 mice; visual discrimination n=320 tracked neurons, 5 mice).

(c) Distribution of orientation preference for the neurons shown in Fig. 6d right, except that values were normalized by the total numbers of neurons imaged n= 1311 and 762 neurons , rather than the number of visually responsive neurons.

Source data are provided as a Source Data file.

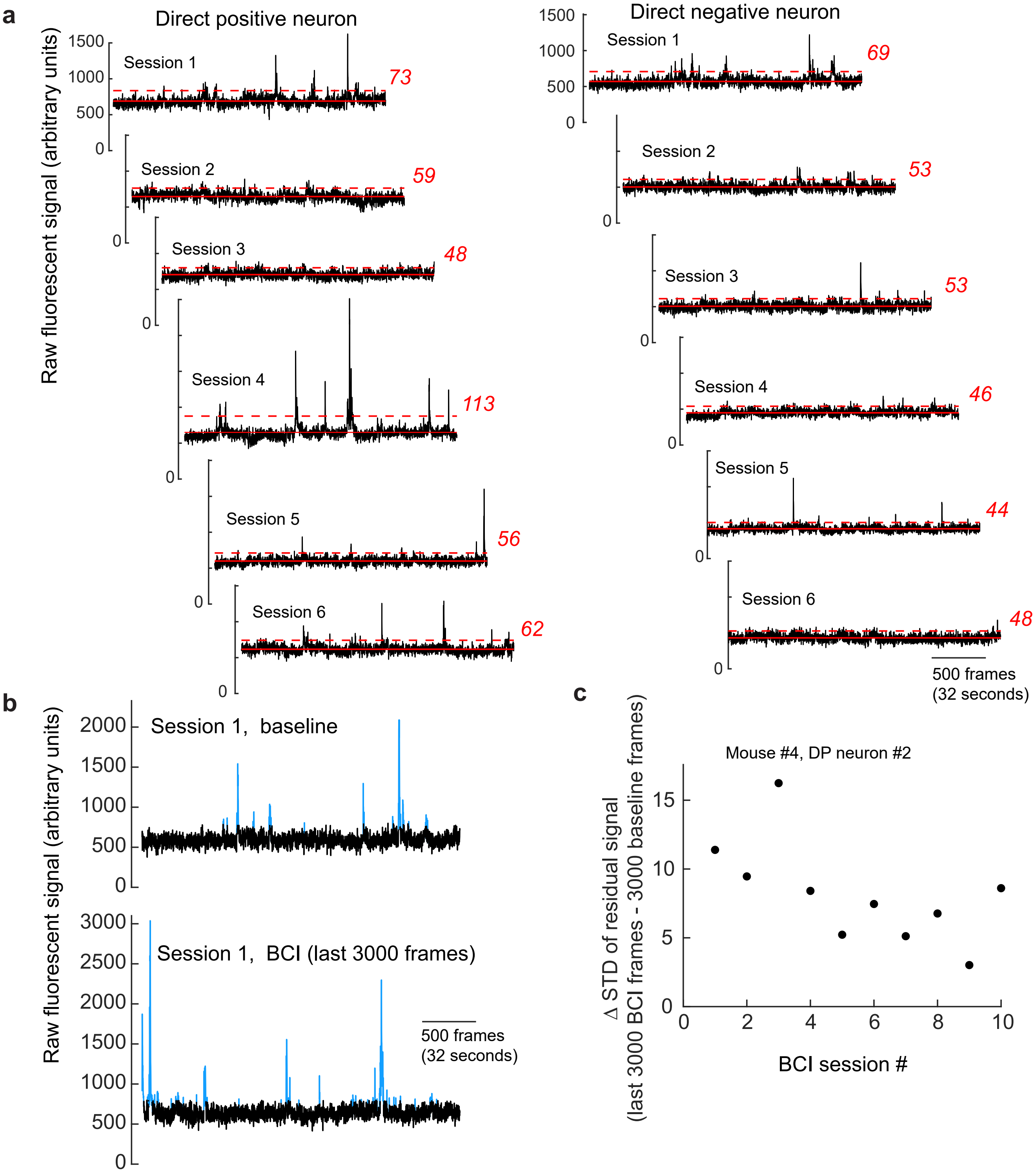

**Supplementary Figure 18. Imaging conditions were stable across and within BCI sessions.**

(a) Example 3-minute spontaneous baseline recorded immediately before the block of BCI trials were initiated, across sessions. The mean fluorescence value across the 3-minute recording is indicated (solid red line), and 1-standard deviation is shown (dashed line and red italicized text). The first session up to the LP session is shown. Data are from mouse #2. Standard deviation was not correlated with session number for any of the direct neurons (Pearson's correlation and p values shown in Supplementary Table 7).

(b) To estimate within-session noise, the change in the standard deviation of the residual signal (black) was calculated, using the 3-minute spontaneous baseline recorded immediately before BCI trials were initiated and the last 3 minutes of the BCI trials for the same session. Calcium events were identified using the 'rmoutliers' function in Matlab and removed (blue) The standard deviation of the residual signal was not correlated for any of the direct neurons (Pearson's correlation and p values shown in Supplementary Table 7). An example of the residual signal is shown for the DP neuron with the median correlation coefficient (mouse #2, DP neuron #3).

(c) The lowest Pearson's correlation p value (mouse #4, DP neuron #2) for the change in the standard deviation of the residual signal versus session number was p= 0.057, n = 10 sessions (Supplementary Table 7). Visual inspection confirmed that there was not a systematic change in the within-session stability of the standard deviation of the residual signal.

Source data are provided as a Source Data file.

| | | | | Experiment | | | | | | |
|---|---|---|---|---|---|---|---|---|---|---|
| Animal ID | Animal label | Sex | Genotype | Auditory task | BCI task | BCI task control | BCI decoding control (Figure 5g) | Visual discrim. task | Visual discrim. control (Figure 6d) | Visual discrim. control (Figure 6h, S17b) |
| slc077_1L | | M | SLC17a7cre | | | | | | | x |
| 2144_1R | mouse 1 | F | EMX1cre | x | x | | | | x | |
| 2317_1R1L | mouse 2 | F | EMX1cre | x | x | | | | | x* |
| 2320_NC | mouse 3 | M | EMX1cre | x | x | | | | | x* |
| 2253_1R | mouse 4 | M | EMX1cre | x | x | | | | | x* |
| 2166_2R | mouse 5 | F | EMX1cre | x | | | | | | |
| slc077_1R | mouse 6 | M | SLC17a7cre | x | | x | | | | |
| slc078_1R | | F | SLC17a7cre | | | | | | | x |
| slc090_1L | mouse 1 | F | SLC17a7cre | | | | | x | | |
| slc091_1L | mouse 2 | M | SLC17a7cre | | | | | x | | |
| slc096_1L | mouse 3 | F | SLC17a7cre | | | | | x | | |
| slc096_1R | mouse 4 | F | SLC17a7cre | | | | | x | | |
| slc096_NC | mouse 5 | F | SLC17a7cre | | | | | x | | |
| slc099_NC | mouse 6 | F | SLC17a7cre | | | | | x | | |
| 2119_NC | | F | EMX1cre | | | | | | x | |
| 2036_2R | | F | EMX1cre | | | | | | x | |
| 2209_2L | | M | EMX1cre | | | | | | x | |
| 2210_NC | | M | EMX1cre | | | | | | x | |
| 2176_1R1L | | M | EMX1cre | | | | | | x | |
| 2452_1R | mouse 1 | M | EMX1cre | | | | x | | | |
| 2452_1R1L | mouse 2 | M | EMX1cre | | | | x | | | |
| 2454_1R | mouse 3 | M | EMX1cre | | | | x | | | |
| 2473_1R | mouse 4 | F | EMX1cre | | | | x | | | |
| 2472_1L | mouse 5 | M | EMX1cre | | | | x | | | |
| 2474_1R1L | mouse 6 | F | EMX1cre | | | | x | | | |

**Supplementary Table 1. Animal sex and genotype information organized by experiment type.**
* The second imaging session occurred after the auditory task training was initiated but before the BCI task training commenced.

| | Direct Neuron ID | | |
|---|---|---|---|
| **Mouse ID** | **DP 1** | **DP2** | **DP3** |
| 1 | -0.039 (5.31E-6) | +0.676 (2.26E-16) | +0.096 (1.92E-05) |
| 2 | -0.411 (3.38E-10 ) | +0.072 (4.79E-2) | -0.140 (6.08E-05) |
| 3 | +0.410 (1.4E-2) | -0.086 (4.60E-2) | -0.086 (2.3E-2) |
| 4 | -0.009 (n.s.) | +0.010 (n.s.) | -0.151 (4.48E-05) |

**Supplementary Table 2. Change in activity at the time of success between session 1 and the LP session for all DP neurons.**

The change in mean activity at the time of success (LP session – session 1) across successful trials is shown; p values of significant K-S tests are in parentheses.

| | | Imaging session | |
|---|---|---|---|
| | **Direct Neuron ID** | **VS baseline 2** | **VS post-learning** |
| Mouse 1 | DP 1 | Not responsive | Not responsive |
| | DP 2 | Not responsive | Not responsive |
| | DP 3 | Not responsive | Not responsive |
| | DN 1 | -0.01 | -0.02 |
| | DN 2 | Not responsive | Not responsive |
| | DN 3 | Not responsive | Not responsive |
| Mouse 2 | DP 1 | -0.02 | -0.03 |
| | DP 2 | -0.02 | -0.03 |
| | DP 3 | Not Tuned | Not responsive |
| | DN 1 | Not responsive | Not responsive |
| | DN 2 | -0.02 | Not responsive |
| | DN 3 | Not Tuned | Not responsive |
| Mouse 3 | DP 1 | -0.05 | -0.03 |
| | DP 2 | Not Responsive | Not responsive |
| | DP 3 | Not Responsive | Not responsive |
| | DN 1 | Not Responsive | Not Responsive |
| | DN 2 | Not Responsive | Not Responsive |
| | DN 3 | Not Responsive | Not Responsive |
| Mouse 4 | DP 1 | -0.04 | -0.03 |
| | DP 2 | -0.03 | -0.03 |
| | DP 3 | -0.03 | -0.04 |
| | DN 1 | -0.03 | -0.03 |
| | DN 2 | -0.03 | 0.02 |
| | DN 3 | -0.03 | Not Responsive |

**Supplementary Table 3. Change in noise correlation values for all direct neurons.**
For a given direct neuron, the median change in pairwise noise correlation for the condition as indicated; all indirect and direct pairs were considered.

| | | Baseline condition | | | | BCI condition | | | |
|---|---|---|---|---|---|---|---|---|---|
| | Direct Neuron ID | Δ Orientation preference (°) | Δ Orientation bandwidth (°) | Δ SF preference (cycles/°) | Δ SF bandwidth (cycles/°) | Δ Orientation preference (°) | Δ Orientation bandwidth (°) | Δ SF preference (cycles/°) | Δ SF bandwidth (cycles/°) |
| Mouse 1 | DP 1 | Not responsive | Not responsive | Not responsive | Not responsive | Not responsive | Not responsive | Not responsive | Not responsive |
| | DP 2 | Not responsive | Not responsive | Not responsive | Not responsive | Not responsive | Not responsive | Not responsive | Not responsive |
| | DP 3 | Not responsive | Not responsive | Not responsive | Not responsive | Not responsive | Not responsive | Not responsive | Not responsive |
| | DN 1 | 9.83 | 29.78 | 0.01 | 0.01 | 72.55 | 32.30 | 0.20 | 0.16 |
| | DN 2 | Not responsive | Not responsive | Not responsive | Not responsive | Not responsive | Not responsive | Not responsive | Not responsive |
| | DN 3 | Not responsive | Not responsive | Not responsive | Not responsive | Not responsive | Not responsive | Not responsive | Not responsive |
| Mouse 2 | DP 1 | 9.54 | 6.63 | 0.00 | 0.00 | 19.54 | 5.63 | 0.01 | 0.00 |
| | DP 2 | 10.86 | 0.85 | 0.00 | 0.00 | 17.41 | 8.30 | 0.00 | 0.00 |
| | DP 3 | Not tuned | Not tuned | Not tuned | Not tuned | Not responsive | Not responsive | Not responsive | Not responsive |
| | DN 1 | Not responsive | Not responsive | Not responsive | Not responsive | Not responsive | Not responsive | Not responsive | Not responsive |
| | DN 2 | 1.81 | 1.13 | 0.02 | 0.01 | Not responsive | Not responsive | Not responsive | Not responsive |
| | DN 3 | Not tuned | Not Tuned | Not tuned | Not Tuned | Not responsive | Not responsive | Not responsive | Not responsive |
| Mouse 3 | DP 1 | 22.00 | 15.36 | 0.06 | 0.06 | 34.27 | 13.28 | 0.06 | 0.02 |
| | DP 2 | Not responsive | Not responsive | Not responsive | Not responsive | Not responsive | Not responsive | Not responsive | Not responsive |
| | DP 3 | Not responsive | Not responsive | Not responsive | Not responsive | Not responsive | Not responsive | Not responsive | Not responsive |
| | DN 1 | Not responsive | Not responsive | Not responsive | Not responsive | Not responsive | Not responsive | Not responsive | Not responsive |
| | DN 2 | Not responsive | Not responsive | Not responsive | Not responsive | Not responsive | Not responsive | Not responsive | Not responsive |
| | DN 3 | Not responsive | Not responsive | Not responsive | Not responsive | Not responsive | Not responsive | Not responsive | Not responsive |
| Mouse 4 | DP 1 | 2.90 | 1.96 | 0.02 | 0.03 | 1.20 | 1.88 | 0.02 | 0.01 |
| | DP 2 | 0.35 | 2.35 | 0.00 | 0.00 | 2.60 | 2.28 | 0.00 | 0.01 |
| | DP 3 | 1.07 | 1.04 | 0.00 | 0.02 | 4.29 | 2.05 | 0.00 | 0.00 |
| | DN 1 | 1.01 | 2.59 | 0.00 | 0.00 | 9.09 | 0.52 | 0.01 | 0.01 |
| | DN 2 | 4.33 | 1.92 | 0.00 | 0.01 | 3.52 | 2.12 | 0.00 | 0.00 |
| | DN 3 | 0.86 | 7.30 | 0.00 | 0.01 | Not responsive | Not responsive | Not responsive | Not responsive |

**Supplementary Table 4. Change in tuning parameters for all direct neurons.**

| | | Baseline condition | | | BCI condition | | |
|---|---|---|---|---|---|---|---|
| | Direct Neuron ID | Median Δ | Minimum Δ | Maximum Δ | Median Δ | Minimum Δ | Maximum Δ |
| Mouse 1 | DP 1 | Not responsive | Not responsive | Not responsive | Not responsive | Not responsive | Not responsive |
| | DP 2 | Not responsive | Not responsive | Not responsive | Not responsive | Not responsive | Not responsive |
| | DP 3 | Not responsive | Not responsive | Not responsive | Not responsive | Not responsive | Not responsive |
| | DN 1 | 0.04 | -0.08 | 0.37 | -0.05 | -0.65 | 0.13 |
| | DN 2 | Not responsive | Not responsive | Not responsive | Not responsive | Not responsive | Not responsive |
| | DN 3 | Not responsive | Not responsive | Not responsive | Not responsive | Not responsive | Not responsive |
| Mouse 2 | DP 1 | 0.01 | -0.91 | 0.56 | 0.00 | -0.47 | 0.60 |
| | DP 2 | 0.00 | -0.11 | 0.45 | 0.00 | -0.21 | 0.40 |
| | DP 3 | Not Tuned | Not Tuned | Not Tuned | Not responsive | Not responsive | Not responsive |
| | DN 1 | Not responsive | Not responsive | Not responsive | Not responsive | Not responsive | Not responsive |
| | DN 2 | 0.03 | -0.24 | 0.16 | Not responsive | Not responsive | Not responsive |
| | DN 3 | Not Tuned | Not Tuned | Not Tuned | Not responsive | Not responsive | Not responsive |
| Mouse 3 | DP 1 | 0.01 | -0.56 | 0.16 | -0.01 | -0.35 | 0.21 |
| | DP 2 | Not Responsive | Not Responsive | Not responsive | Not responsive | Not responsive | Not responsive |
| | DP 3 | Not Responsive | Not Responsive | Not responsive | Not responsive | Not responsive | Not responsive |
| | DN 1 | Not Responsive | Not Responsive | Not responsive | Not Responsive | Not Responsive | Not responsive |
| | DN 2 | Not Responsive | Not Responsive | Not responsive | Not Responsive | Not Responsive | Not responsive |
| | DN 3 | Not Responsive | Not Responsive | Not responsive | Not Responsive | Not Responsive | Not responsive |
| Mouse 4 | DP 1 | 0.00 | -0.22 | 0.31 | 0.03 | -0.23 | 0.07 |
| | DP 2 | 0.02 | -0.06 | 0.16 | 0.00 | -0.05 | 0.11 |
| | DP 3 | -0.01 | -0.28 | 0.06 | -0.01 | -0.29 | 0.08 |
| | DN 1 | 0.00 | -0.16 | 0.25 | 0.01 | -0.33 | 0.34 |
| | DN 2 | 0.00 | -0.29 | 0.13 | 0.00 | -0.18 | 0.25 |
| | DN 3 | 0.00 | -0.26 | 0.42 | Not Responsive | Not Responsive | Not Responsive |

**Supplementary Table 5. Change in signal correlation values for all direct neurons.**
For a given direct neuron, the median change in pairwise signal correlation for the condition as indicated; all indirect and direct pairs were considered.  In addition, the minimum and maximum changes are included.

| | Direct Neuron ID | Baseline condition | BCI condition |
|---|---|---|---|
| | | Δ amplitude (deconvolved events) | |
| Mouse 1 | DP 1 | Not responsive | Not responsive |
| | DP 2 | Not responsive | Not responsive |
| | DP 3 | Not responsive | Not responsive |
| | DN 1 | 7.38 | 11.57 |
| | DN 2 | Not responsive | Not responsive |
| | DN 3 | Not responsive | Not segmented |
| Mouse 2 | DP 1 | 14.07 | 5.00 |
| | DP 2 | 8.38 | 14.18 |
| | DP 3 | Not Tuned | Not responsive |
| | DN 1 | Not responsive | Not responsive |
| | DN 2 | 3.27 | Not responsive |
| | DN 3 | Not Tuned | Not responsive |
| Mouse 3 | DP 1 | 5.50 | 0.83 |
| | DP 2 | Not responsive | Not responsive |
| | DP 3 | Not responsive | Not responsive |
| | DN 1 | Not responsive | Not responsive |
| | DN 2 | Not responsive | Not responsive |
| | DN 3 | Not responsive | Not responsive |
| Mouse 4 | DP 1 | 2.37 | 3.74 |
| | DP 2 | 4.94 | 4.54 |
| | DP 3 | 2.10 | 0.42 |
| | DN 1 | 3.40 | 1.77 |
| | DN 2 | 5.66 | 1.34 |
| | DN 3 | 0.50 | Not responsive |

**Supplementary Table 6. Change in the amplitude of the preferred stimulus response for all direct neurons.**

| Pearson's correlation of baseline standard devidation and session number | | | | | | |
|---|---|---|---|---|---|---|
| p values | | | | | | |
| Mouse ID | DP1 | DP2 | DP3 | DN1 | DN2 | DN3 |
| 1 | 0.383 | 0.991 | 0.15 | 0.046 | 0.022 | 0.015 |
| 2 | 0.068 | 0.98 | 0.51 | 0.05 | 0.538 | 0.845 |
| 3 | 0.726 | 0.85 | 0.599 | 0.778 | 0.294 | 0.24 |
| 4 | 0.494 | 0.799 | 0.241 | 0.528 | 0.532 | 0.107 |
| r values | | | | | | |
| Mouse ID | DP1 | DP2 | DP3 | DN1 | DN2 | DN3 |
| 1 | 0.310 | -0.004 | 0.491 | -0.641 | -0.707 | -0.736 |
| 2 | -0.780 | 0.013 | 0.340 | -0.812 | -0.319 | -0.104 |
| 3 | 0.148 | 0.080 | 0.221 | 0.119 | -0.425 | 0.470 |
| 4 | 0.246 | 0.093 | 0.409 | 0.227 | 0.225 | 0.541 |

| Pearson's correlation of the within-session change in the standard deviation of the residual signal and session number | | | | | | |
|---|---|---|---|---|---|---|
| p values | | | | | | |
| Mouse ID | DP1 | DP2 | DP3 | DN1 | DN2 | DN3 |
| 1 | 0.132 | 0.851 | 0.534 | 0.171 | 0.176 | 0.504 |
| 2 | 0.795 | 0.401 | 0.653 | 0.774 | 0.694 | 0.796 |
| 3 | 0.859 | 0.151 | 0.903 | 0.560 | 0.193 | 0.116 |
| 4 | 0.991 | 0.057* | 0.689 | 0.091 | 0.675 | 0.600 |
| r values | | | | | | |
| Mouse ID | DP1 | DP2 | DP3 | DN1 | DN2 | DN3 |
| 1 | -0.510 | 0.068 | -0.224 | -0.470 | -0.465 | -0.240 |
| 2 | -0.138 | 0.425 | -0.236 | -0.152 | -0.207 | 0.137 |
| 3 | -0.076 | -0.557 | 0.052 | -0.244 | -0.514 | -0.600 |
| 4 | -0.004 | -0.618 | -0.145 | -0.562 | -0.152 | 0.189 |

**Supplementary Table 7. Cross-session and within session stability of raw calcium signal for all direct neurons.**

Note, the p value of one direction neuron (mouse #4, DP neuron #2) was low, p= 0.057. Visual inspection of the data confirmed that there was not a systematic change in the within-session stability of the standard deviation of the residual signal Supplementary Fig. 18c).
